# Mendelian randomisation analysis of the effect of educational attainment and cognitive ability on smoking behaviour

**DOI:** 10.1101/299826

**Authors:** Eleanor Sanderson, George Davey Smith, Jack Bowden, Marcus R. Munafò

## Abstract

Recent analyses have shown educational attainment to be associated with a number of health outcomes. This association may, in part, be due to an effect of educational attainment on smoking behaviour. In this study we apply a multivariable Mendelian randomisation design to determine whether the effect of educational attainment on smoking behaviour could be due to educational attainment or general cognitive ability. We use individual data from the UK Biobank study (N = 120,050) and summary data from large GWAS studies of educational attainment, cognitive ability and smoking behaviour. Our results show that more years of education are associated with a reduced likelihood of smoking which is not due to an effect of general cognitive ability on smoking behaviour. Given the considerable physical harms associated with smoking, the effect of educational attainment on smoking is likely to contribute to the health inequalities associated with differences in educational attainment.

## Introduction

Smoking remains the largest single, modifiable contributor to morbidity and premature mortality in high income countries such as the United Kingdom.^1^ However, while smoking prevalence has declined steadily since the 1950s, this overall decline masks considerable differences across society. Specifically, while smoking prevalence has declined substantially in higher socioeconomic groups, the decline has been considerably less in lower socioeconomic groups, so that smoking is now strongly socially patterned and an important determinant of health inequalities.

Recent analyses have suggested that educational attainment is an important determinant of health outcomes such as coronary heart disease (CHD). Using genetic variants associated with educational attainment in Mendelian randomisation (MR) analyses,^2^ Tillmann and colleagues provided evidence that lower educational attainment is causally related to increased risk of CHD.^3^ Using similar methods, Gage and colleagues showed similar effects with respect to a range of smoking behaviours, with lower educational attainment related to increased risk of smoking initiation and, if a smoker, increased heaviness of smoking and reduced likelihood of smoking cessation^4^. Carter and colleagues show that smoking mediates the observed effect of education on cardiovascular disease.^5^ It is therefore plausible that some of the effect of educational attainment on risk of CHD (and other adverse health outcomes) is mediated via effects on smoking behaviour.

However, one limitation of analyses of educational attainment is that they do not allow us to distinguish between the effects of education per se, versus the effects of general cognitive ability (which is strongly associated with higher educational attainment).^6–8^ This distinction has important implications when understanding the potential role of education in reducing health inequalities – if apparent protective effects of educational attainment are in fact due to general cognitive ability, then increasing years in education will not necessarily improve health outcomes or reduce health inequalities. In other words, elucidating the independent effects of educational attainment versus general cognitive ability is critical to understanding the nature of health inequalities associated with educational status.

In this study we used genetic variants associated with educational attainment, together with genetic variants associated with general cognitive ability, in a multivariable MR framework using individual level data to determine the unique effects of each on smoking behaviour. This method, described by Sanderson and colleagues^9^ allows multiple genetic instruments, capturing distinct exposures, to be investigated simultaneously (even when these exposures are highly correlated, as in the case of educational attainment and general cognitive ability). We show that the effect of educational attainment we observed in our univariable MR study is also observed in our multivariable MR analysis. However, the effect of cognitive ability on smoking behaviour observed in our univariable MR analysis attenuates when educational attainment is also included in the multivariable MR analysis. From this we conclude that the effect of educational attainment on smoking behaviour is not due to an effect of general cognitive ability on smoking behaviour and the effect of general cognitive ability on smoking observed in the univariable MR analysis is acting via the effect of educational attainment on smoking behaviour.

## Results

### Observational analysis

Observational analyses, controlling for the full set of controls using ordinary least squares (OLS) regression, indicated a small negative association between educational attainment, general cognitive ability and current smoking. These results also indicated a negative association between education and smoking initiation, and a positive association between general cognitive ability and smoking initiation. Educational attainment and general cognitive ability were both positively associated with smoking cessation. These results are shown in Tables 1–3.

**Table 1.** The effect of education and general cognitive ability on current smoking.

|  | OLS | Single<br>variable MR | Single<br>variable MR | Multivariable<br>MR |
| --- | --- | --- | --- | --- |
| ***Age completed education*** |  |  |  |  |
| Effect | -0.007 | -0.027 |  | -0.041 |
| SE | 0.0004 | 0.005 |  | 0.012 |
| 95% CI | -0.008, -0.007 | -0.037, -0.017 |  | -0.064, -0.018 |
| P-value | <0.001 | <0.001 |  | 0.001 |
| F | - | 556.17 |  | 483.19 |
| Conditional F | - | - |  | 85.89 |
| ***Cognitive ability score*** |  |  |  |  |
| Effect | -0.008 |  | -0.026 | 0.047 |
| SE | 0.001 |  | 0.009 | 0.027 |
| 95% CI | -0.010, -0.007 |  | -0.044, -0.008 | -0.005, 0.100 |
| P-value | <0.001 |  | 0.005 | 0.076 |
| F | - |  | 887.78 | 544.24 |
| Conditional F | - |  | - | 86.78 |

Estimates of the effect of education and general cognitive ability on current smoking from OLS, multivariable MR and single variable MR regressions from analysis of individual level data. Estimates given are risk difference effect estimates. All regressions also include a full set of adjustments: age, sex, year of birth and gender interacted with year of birth. Mendelian randomisation regressions are also adjusted for 40 genetic principal components. Non-European and related individuals have been excluded from the analysis. Total sample size 120,050.

**Table 2.**
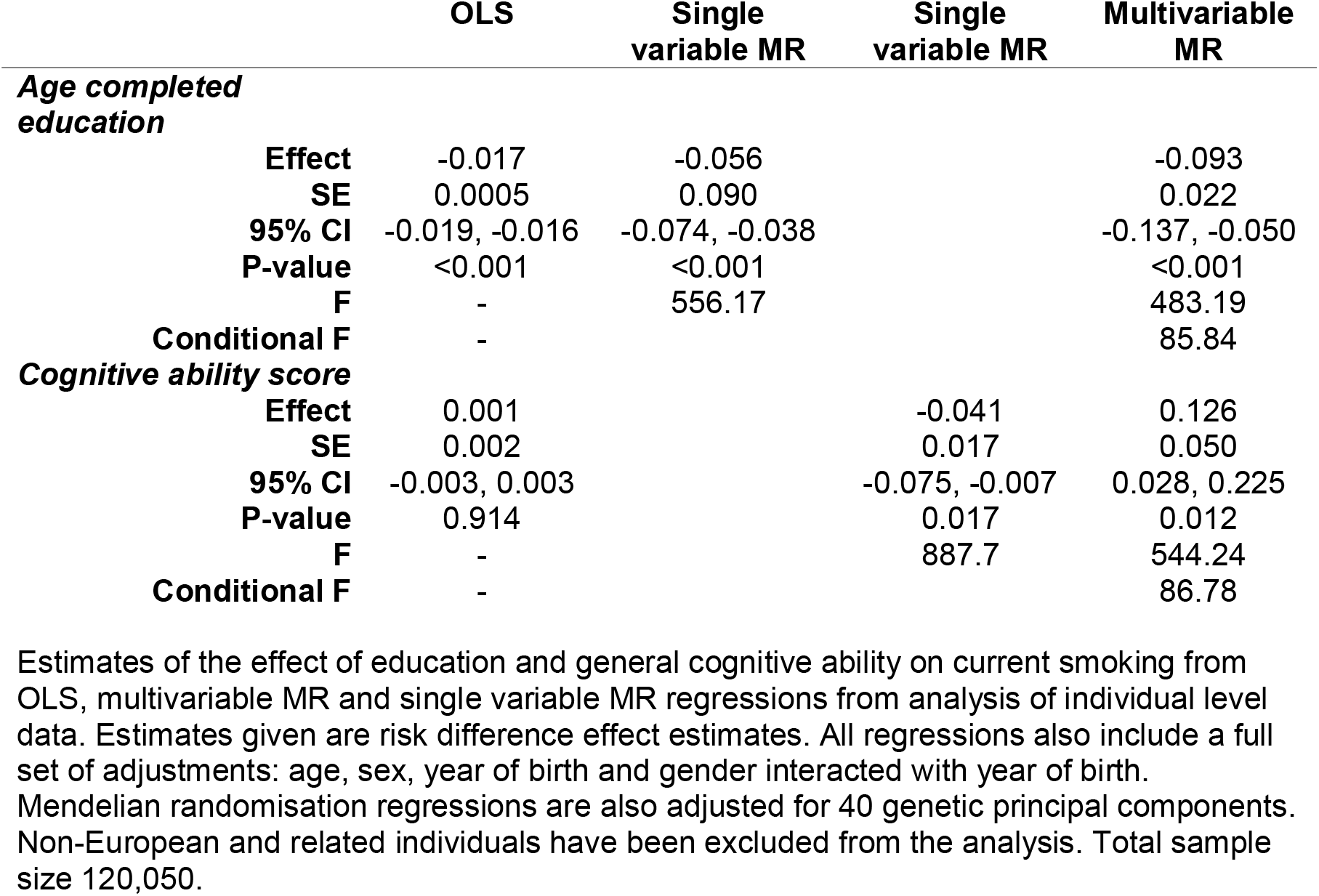
The effect of education and general cognitive ability on smoking initiation

**Table 3.**
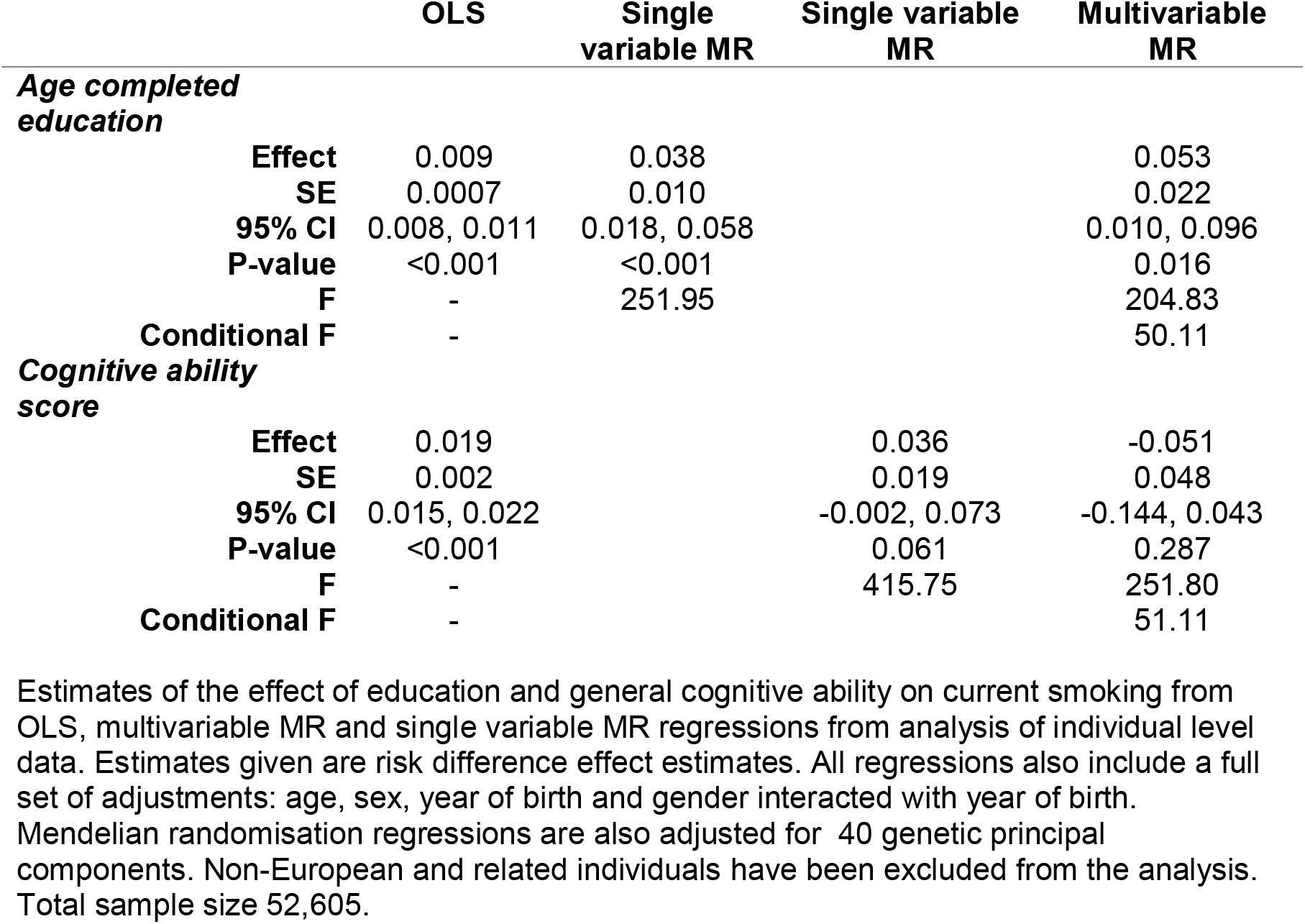
The effect of education and general cognitive ability on smoking cessation.

### Individual-level analysis

In all of these analyses we used one instrument in each of the univariable MR analyses and two instruments in the multivariable MR analysis. These instruments are risk scores for each exposure created using SNPs that have been shown to be associated with the exposure. The F-statistics, given in Tables 1 – 3, show that the genetic instruments used strongly predict education and general cognitive ability. In all of our multivariable MR estimations the conditional F-statistic is larger than the conventional value of 10 used to indicate a strong instrument, and so the instruments used are strong enough to uniquely predict both of the exposure variables in the multivariable MR analyses.

The unconditional univariable MR results indicate a negative effect of education on current smoking with each year of education leading to a 2.6% decrease in the probability of being a smoker. One standard deviation in age completed education is 2.4 years; therefore this corresponds to a standard deviation increase in education leading to a 6.2% decrease in the probability of being a smoker. Similar results are seen for the direct, conditional, effect estimated by multivariable MR with each additional year of education leading to a 4.1% decrease in the probability of being a current smoker, corresponding to an increase in education of one standard deviation leading to a 9.8% decrease in the probability of being a current smoker.

The unconditional effect of general cognitive ability estimates a negative effect of increased general cognitive ability on current smoking with a standard deviation increase in general cognitive ability leading to a decrease in current smoking of 2.6%. The estimates for multivariable MR differed substantially from the single variable MR result, there was a high level of uncertainty around the estimated effect with a 95% confidence interval of a 0.5% decrease to a 10.2% increase and no evidence of an effect of general cognitive ability on smoking behaviour. One explanation for the difference observed between the univariable and multivariable MR estimates for the effect of general cognitive ability on smoking is that general cognitive ability affects smoking through its effect on educational attainment, rather than through a direct effect on smoking, i.e. the relationship illustrated in Fig 2.

**Figure 1:**
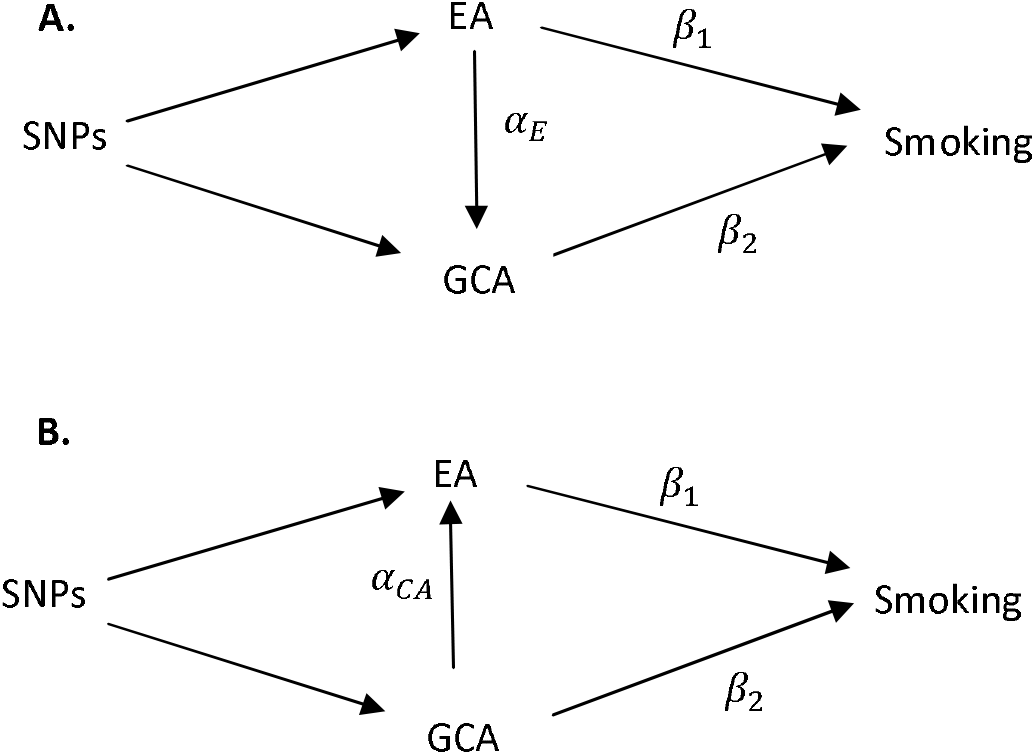
Potential relationships between education, cognitive ability and smoking. Potential Directed Acyclic Graphs (DAG) for the relationship between educational attainment, general cognitive ability and smoking behaviour. A. general cognitive ability is a mediator of some of the relationship between educational attainment and smoking. B. educational attainment is a mediator of some of the relationship between general cognitive ability and smoking.

**Figure 2:**
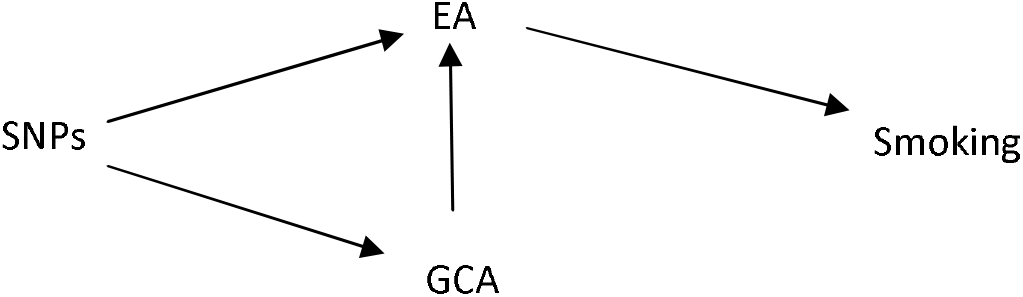
Relationship between education, cognitive ability and smoking suggested by the results obtained. DAG for the relationship between educational attainment, general cognitive ability and smoking suggested by the results obtained.

The results for smoking initiation showed a similar pattern to the results for being a current smoker. The univariable MR results showed that increases in both education and general cognitive ability lead to decreases in the probability of ever having been a smoker. However the multivariable MR results show a negative direct effect of education on smoking and a positive direct effect of general cognitive ability on smoking. An increase in education of one year causes a 9.1% decrease in the probability of having ever smoked, this is equivalent to a 21.8% decrease in the probability of ever smoking for a standard deviation increase in years of education. An increase in general cognitive ability of one standard deviation causes a 12.5% increase in the probability of ever having smoked. There is however still a large level of uncertainty around this result with a 95% confidence interval of a 2.5% increase to a 22.5% increase.

The results for being a former smoker showed that higher education leads to a higher probability of having quit smoking, but did not indicate a direct effect of general cognitive ability on quitting smoking. These results reflect the pattern seen in the results for currently or ever smoking showing that increased education makes individuals more likely to quit smoking once they have started as well as being less likely to start smoking in the first place.

As a sensitivity analysis we repeated the UK Biobank analyses with self-reported age at which individual completed education. Individuals who had not completed a degree were also asked to report the age at which they left school. The distribution of the age individuals left school for each educational qualification is given in Supplementary Table 1. As some individuals only completed the cognition test via an online test we additionally repeated the analysis using only the cognitive ability scores completed at the clinic. Results from these analyses are given in Supplementary Tables 2 – 3. Neither of these analyses give results that are substantially different from those presented here.

UK Biobank has been shown to be unrepresentative of the UK population with individuals in UK Biobank reporting higher socioeconomic status and lower rates of smoking than the general population.^10^ If participation in the study is influenced by educational attainment, cognitive ability or smoking this may cause selection bias in our estimation results.^11,12^ The level of this bias is unknown and limited methods are currently available to correct for selection bias of this type. Following Hughes and colleagues,^11^ we re-estimate our results with an education weighting based on the proportion of individuals in the population of the same age range as the UK Biobank sample who report leaving school before age 16 or who have obtained a degree in the 2011 UK census. These results are given in Supplementary Table 4 and show that our estimation results are not substantially influenced by the weighting.

### Summary-level analyses

Each additional year of education in the univariable MR analysis was associated with a 35% decrease in the probability of starting smoking. Once the effect of cognitive ability had been controlled for in the multivariable MR analysis each additional year of educational attainment was associated with a 40% decrease in the probability of smoking. However, there is limited evidence from these results that cognitive ability is associated with a change in the rate of smoking initiation. These results are given in Table 4.

**Table 4.**
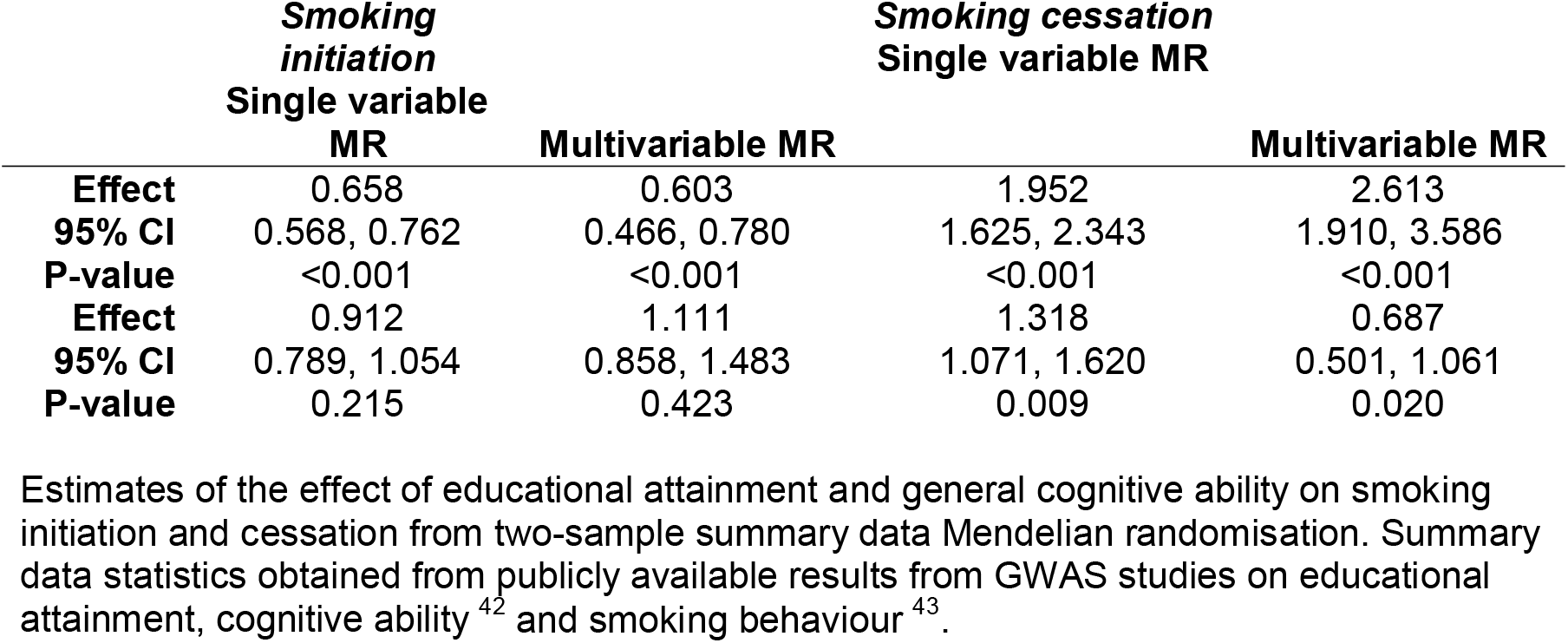
Estimates of the effect of educational attainment and general cognitive ability; Odds ratio estimates from summary level data analysis.

The summary data analysis showed that each additional year of education makes individuals who have taken up smoking twice as likely to have quit in the univariable MR analysis and 2.6 times as likely to quit smoking in the multivariable MR analysis once the effect of cognitive ability has been controlled for. There was also evidence that higher cognitive ability is associated with being more likely to quit smoking. However, this association disappears when educational attainment is controlled for in the multivariable MR analysis. These results are given in Table 4.

These results support the findings from the individual-level analysis and the conclusion that there is a direct effect of educational attainment on smoking behaviour and limited direct effect of cognitive ability on smoking behaviour. The size of the estimated effects in the individual level and summary data analysis are not directly comparable as the individual level analysis estimates the effect of education and cognitive function on smoking at a particular point in time whereas the summary data analysis will represent an estimate of the lifetime effect on smoking behaviour.

Horizontal pleiotropy, where the SNPs used in our MR study have an effect on smoking which is not through their effect on educational attainment or cognitive ability introduces bias into MR estimation results.^13^ High levels of heterogeneity in the estimated effects from each SNP are an indication of potential pleiotropic effects of some of the SNPs associated with educational attainment or general cognitive ability used in the analysis on smoking behaviour. To test for potential pleiotropy we estimate modified Cochran Q statistic^9,14^ for each of our summary data multivariable MR models, reported in Supplementary Table 5. Supplementary Figure 1 shows the individual SNP Q statistics which give the contribution of each SNP to the overall Q statistic. We re-estimate our summary data MR analysis excluding any SNPs with individual Q statistics with a p-value less than 0.05 to exclude SNPs with a potentially pleiotropic effect. This results in 32 SNPs being removed from the analysis of smoking initiation and 17 SNPs being removed from the analysis of smoking cessation. The results from this re-estimation are given in Supplementary Table 6, they show that removing these results makes no substantive difference to our results.

Under a set of assumptions (known as the InSIDE assumptions) MR Egger can detect directional pleiotropy and give causal effect estimates that are robust to such pleiotropy.^14^ This method has been extended to two-sample summary data multivariable MR analysis^15^ and therefore we estimate MR Egger for both our univariable and multivariable summary data MR estimates. These results are given in Supplementary Table 7. The estimate of the constant in an MR Egger regression is a measure of the level of directional pleiotropy in the analysis. In all of the analyses the constant is estimated to be close to zero, with narrow 95% confidence intervals around zero. These results, combined with those from our analysis of the Q statistic, suggest that the estimation results are not substantially affected by horizontal pleiotropy through the SNPs affecting the outcome via an effect other than their effect on educational attainment and general cognitive ability.

## Discussion

The univariable MR estimates give the total effect of a change in education and general cognitive ability on three smoking phenotypes including any effect of general cognitive ability on education and vice versa. Our multivariable MR results allow us to estimate the direct effects of each of education and general cognitive ability on smoking behaviour conditional on general cognitive ability and education respectively. Overall, our results suggest that more years of education leads to a reduced likelihood of smoking and, among those who do smoke, a greater likelihood of cessation. Given the considerable physical harms associated with smoking, this is likely to account for a substantial proportion of health inequalities associated with differences in educational attainment. The similarity between the unconditional and conditional effects of educational attainment on all aspects of smoking behaviour considered, suggests that education has an effect on smoking behaviour. However, the large difference between the univariable and multivariable MR estimates for the effect of general cognitive ability on smoking suggests that a large part of the total effect of general cognitive ability on smoking behaviour observed in the conventional MR analysis is due to general cognitive ability affecting smoking through an effect on educational attainment rather than a direct effect of general cognitive ability on smoking behaviour. These results are consistent with recent findings reported by Davies and colleagues, who used the natural experiment of an increase in school leaving age to show that remaining in school causally reduces the risk of adverse health outcomes, including likelihood of smoking^16^ and with observational studies which have shown an attenuation in the effect of cognitive ability on the likelihood of smoking once educational attainment is controlled for.^17,18^

Estimating effects in this way, using MR methods, does not necessarily clarify the mechanistic pathways that link exposures and outcomes. Our results highlight an important, unintended, public health benefit of education and support a causal pathway from educational attainment to smoking behaviour that appears to be independent of general cognitive ability. These results suggest that the observed association between education and smoking behaviour is not due to a direct effect of general cognitive ability on smoking behaviour. However, these results do not identify the specific mechanism or aspect of the current educational system by which the effect of education on smoking occurs and the pathway is likely to be complex, meaning further work is required to understand the particular mechanisms underlying this finding. More years in education might influence smoking via greater awareness of the harms of smoking, increased self-efficacy, exposure to social groups where smoking is less common, and so on. While years in education might be considered a target for intervention, there is only limited scope to extend this. Identifying other links in the pathway may also reveal other putative intervention targets that are potentially more tractable, providing an additional public health benefit. This should be the subject of future research, and in part will depend on the identification of genetic variants associated with, for example, self-efficacy beliefs if MR methods are to be applied.

There are a number of important limitations to this study that should be considered when interpreting these results. First, Mendelian randomization is not without limitations and, critically, relies on key assumptions, in particular regarding the validity of the genetic variants used as instruments. We have shown that the genetic variants we used are strongly associated with both education and general cognitive ability. We also assessed the robustness of our analyses to potential pleiotropy through sensitivity analysis using methods that rely on different assumptions (e.g., MR Egger and outlier removal)^19^, and these produced similar results. Second, we cannot rule out the possibility of dynastic effects – offspring will share on average 50% of the variegated genotype of their parents. Parental genotype will affect parental education levels, which will therefore be correlated with offspring genotype. Parental genotype has previously been shown to be correlated with offspring educational attainment.^20^ If parental education has an effect on offspring smoking that does not operate through offspring education this would represent potential violation of assumption IV2, and the effects we observed may in fact be due to effects of parental education on the offspring environment that have an impact on smoking behaviour. This relationship is illustrated in Fig. 3. While we cannot rule out this possibility, our sensitivity analyses do not show evidence of a violation of assumption IV2. Crucially, other studies which have estimated the effect of educational attainment on smoking behaviour using alternative methods such as structural modelling^21^, sibling analyses^22^, or alternative IV strategies (using a natural experiment)^16^ have found similar effects of education on smoking behaviour to those we observe. The different design of each of these studies means that they are subject to *different* potential sources of bias; particularly neither the study using sibling analysis, nor the natural experiment IV study, which took advantage of a policy change in the compulsory school leaving age, are subject to potential bias from dynastic effects. Our results add to this body of evidence; while each study has limitations, comparing the set of results within a triangulation framework^23^ strengthens our ability to draw the conclusion that increasing education leads to a decrease in smoking. The potential for dynastic effects and parental nurturing to bias our results could be avoided by repeating our analysis using within family designs such as variations within sets of siblings who will be subject to the same dynastic effects. Such a study would be subject to a different set of assumptions and potential biases to those in our study and the studies described above and would contribute further to the body of evidence. UK Biobank contains a number of sibling pairs, however as the study was not recruited as a family design the total number is small. Specifically, there are 11,448 individuals from 5,613 families with at least one sibling in UK Biobank who pass all other data restrictions for our analysis. Although we would ideally repeat our analysis using a family design in this sample to obtain estimated effects free from any potential dynastic effects this sample is not large enough to obtain a result. This is shown by the power calculations given in Supplementary Table 8 which show that with a sample size of 11,448 individuals from 5,724 sibling pairs the power to detect an effect of educational attainment on smoking behaviour similar in magnitude to those we observe in our main analysis with this sample is low. Therefore this analysis remains an area for future research. Third, the UK Biobank is highly unrepresentative of the general UK population, and this selection bias may introduce biases from which MR methods are not immune.^24^ For example, if educational attainment and smoking behaviour are both associated with participation in UK Biobank, then restricting our analyses to this sample may introduce collider bias, and bias associations between these variables (or even introduce spurious associations between them). However, if individuals are more likely to participate if they are more educated and are less likely to participate if they smoke, as has been observed,^10^ this will decrease the association between education and smoking observed. Alternatively, if participation is only determined by education then, if the association between education and smoking is the same in the whole population as in the sample, this selection will have no effect on the results obtained. Our sensitivity analyses attempt to correct for the selection on education by reweighting the analysis however it will be important to replicate these findings in other samples that are either more representative of the general population or are subject to different selection biases. Fourth, the interpretation of multivariable MR results is nuanced and potentially complex. Specifically, the estimate for educational attainment is adjusted for general cognitive ability and vice versa. This means that the estimated effect of education is the effect given a constant level of general cognitive ability, and the effect for general cognitive ability is the effect given a constant level of education. While the interpretation of our educational attainment results is relatively straightforward, the interpretation of our conditional general cognitive ability results (which imply increased likelihood of smoking with increasing general cognitive ability) is not in line with what we might expect. As general cognitive ability affects education what this (weak) effect may reflect is the fact that individuals with high levels of general cognitive ability who do not achieve the educational attainment that might be expected are likely to have done so for specific reasons that are themselves likely to influence smoking. This result indicates that the beneficial effect of general cognitive ability on smoking behaviour observed in the conventional MR analysis is acting via the effect of general cognitive ability on education, and the consequent effect of education on smoking behaviour. This suggests that an increase in general cognitive ability that was not associated with a corresponding increase in educational attainment would have a limited effect on smoking behaviour.

**Figure 3:**
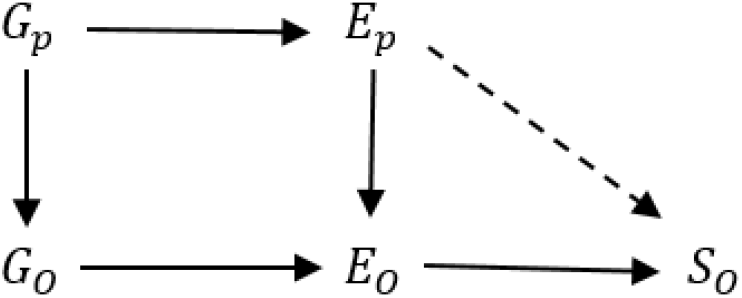
Potential for bias due to dynastic effects. DAG showing how dynastic effects can introduce bias into an MR model. *G_p_* is parental genotype, *E_p_* parental educational attainment, *G_O_* is offspring genotype, *E_O_* offspring educational attainment and *S_O_* offspring smoking. The dashed line represents the relationship required for dynastic effects to introduce bias in an MR study.

In summary, our results indicate that the observed association between educational attainment and smoking is unlikely to be due to effects of general cognitive ability on educational attainment and smoking behaviour. This suggests that a part of the health inequalities associated with differences in educational outcomes may be due to smoking, and that education represents a potential target for intervention to reduce health inequalities. Future research may identify other putative targets on the causal pathway from educational attainment and smoking behaviour and should also explore whether there are similar relationships to other health behaviours such as alcohol consumption and diet.

## Methods

### Individual level data

For our individual level analysis we used data from UK Biobank. This is a population-based health research resource consisting of approximately 500,000 people, aged between 38 years and 73 years, who were recruited between the years 2006 and 2010 from across the UK.^25^ Participants provided a range of information (such as demographics, health status, lifestyle measures, cognitive testing, personality self-report, and physical and mental health measures) via questionnaires and interviews. A full description of the study design, participants and quality control (QC) methods is given by Collins (2012).^26^ Informed consent was obtained from study participants by UK Biobank. UK Biobank received ethics approval from the Research Ethics Committee (REC reference for UK Biobank is 11/NW/0382). Individuals with sex-mismatch (derived by comparing genetic sex and reported sex) or individuals with sex chromosome aneuploidy were excluded from the analysis (n=848). We restricted the sample to individuals of European ancestry who have very similar ancestral backgrounds according to the PCA (n=423,166), as described by Bycroft.^27^ Estimated kinship coefficients using the KING toolset^28^ identified 107,162 pairs of related individuals in the dataset.^27^ In order to mitigate this feature in the data a standard algorithm was then applied to preferentially remove the most highly related individuals in our sample (71,133). Full details are given elsewhere.^29^

The SNPs that we used to create our instrument for general cognitive ability were discovered using the interim release of UK Biobank, and so we additionally excluded all individuals who were included in the interim release (94,985). This gives a resulting potential sample size of 257,048 individuals. Of these individuals we only included those with observations for all of the exposures and outcomes included in the analysis. A total of 127,606 of the individuals eligible to be included in our sample completed the general cognitive ability questions. However, 7,564 of these individuals had missing data for one of the other exposures or controls included in the analysis, giving a final sample size of 120,050 individuals.

Individuals in UK Biobank were asked to report their highest educational qualification. For each of the levels of education a corresponding age at which the individual would have completed their education has been assigned. These are described in Table 5.

**Table 5.** Highest educational qualifications.

| Highest Educational qualification | Age completed education | % of main sample |
| --- | --- | --- |
| None | 15 | 12.1 |
| CSE / O level / GCSE | 16 | 26.8 |
| NVQ / HND / HNC | 18 | 6.0 |
| A level | 18 | 12.4 |
| Other professional qualification * | 20 | 5.1 |
| College or University degree | 21 | 37.5 |

CSE / O level / GCSE: Academic qualification at 16+; NVQ / HND / HNC: Vocational qualification; A level: Academic qualification at 18+; \* e.g. nursing/teaching etc.

General cognitive ability was defined as the results from a ‘verbal-numeric reasoning’ test, labelled fluid-intelligence in UK Biobank, measured as the number of correct answers recorded in a series of 13 questions designed to capture verbal and arithmetical ability.^30^ This test has been shown to correlate highly with alternative measures of cognitive ability.^31,32^ The test was completed by participants during a questionnaire as part of their initial clinic visit or through an online questionnaire completed after the visit. The score is the number of questions correctly completed by the individuals in the time allowed and ranges between 0 and 13 with a mean of 6.0 and a standard deviation of 2.1. For the analyses reported here, the score has been standardised to have a mean of 0 and a standard deviation of 1.

All participants were asked if they currently smoked and if they had ever smoked. From this information we constructed three variables: 1) a binary variable for current smokers/current non-smokers; 2) a binary smoking initiation variable for ever/never smokers; and 3) among individuals who have ever smoked, a smoking cessation variable for former vs current smokers. The total sample size is 120,050 of which 9,252 (7.7%) are current smokers and 52,605 (43.8%) are ever smokers. Of the ever smokers 43,353 (82.4%) are former smokers.

A recent GWAS of educational attainment by Okbay and colleagues identified 74 SNPs at genome-wide significance that associated with years of education completed.^33^ A polygenic risk score for education was created for each of the individuals in our sample using the genome-wide significant SNPs from this GWAS. Each individual’s risk score was calculated as the weighted total number of education increasing alleles they had for these SNPs, weighted by the size of the effect of the SNP in the original GWAS. Three of the SNPs from this GWAS where not available in the UK Biobank data and were substituted; all of the substitutes used where in perfect LD with the original SNP.

For general cognitive ability we created a polygenic risk score for each individual using the results from a recent GWAS of general cognitive ability by Sniekers and colleagues, which identified 18 SNPs at genome-wide significance.^34^ Again, the risk score was calculated as the weighted number of general cognitive ability increasing alleles each individual had for these SNPs, weighted by the effect of the SNP in the GWAS.

Each polygenic risk score was created using the association observed between each SNPs and the exposure in a sample that did not include our main analysis sample. This means that these SNPs can be used as potentially valid instruments for our analysis in UK Biobank (with the interim release excluded). The importance of using SNPs discovered in samples that do not include the analysis sample is discussed elsewhere.^35,36^

We use Mendelian randomisation (MR) analyses to estimate the effect of educational attainment and general cognitive ability on smoking behaviour.^37,38^ MR utilises the random allocation of genetic variants at conception to understand the causal effect of a phenotype on an outcome. This random allocation of genetic variants means that they will be unrelated to other traits such as lifestyle and socioeconomic position that bias the observational association between educational attainment, general cognitive ability and smoking behaviour and so are potentially valid instruments in an instrumental variable analysis.^37,38^ For the genetic variants to be valid instruments they must satisfy three instrumental variable assumptions; they must be

IV1. Associated with at least one of the exposures,
IV2. Independent of any factors confounding the relationship between the exposures and the outcome;
IV3. Independent of the outcome given the exposures and the confounders.

We conducted univariable MR analysis of individual level data for each of the exposures on each of our three outcomes using the SNP score for education as the instrument for educational attainment and the SNP score for general cognitive ability as the instrument for general cognitive ability. These results give the unconditional effect of each of education and general cognitive ability on smoking, i.e. the total effect of education and general cognitive ability on each of the smoking phenotypes, including any effect of education on general cognitive ability and any effect of general cognitive ability on education. These effects are illustrated in Fig. 1. If general cognitive ability mediates the relationship between educational attainment and smoking behaviour *α_E_* ≠ 0 and the total effect of education on smoking is given by *β*_1_ *α_E_β*_2_ (Fig. 1A). However, if educational attainment mediates the relationship between general cognitive ability and smoking behaviour *α_CA_* ≠ 0 and the total effect of general cognitive ability on smoking is given by *β*_2_ + *α_CA_β*_1_ (Fig. 1B). These results are susceptible to horizontal pleiotropy, where the genetic instruments are associated with a confounder of the relationship between the exposure and outcome and so have an effect on the outcome that doesn’t operate through the exposure, a violation of the assumption IV2, as any direct effect of the education score on general cognitive ability or the general cognitive ability score on educational attainment will affect the results obtained from this analysis.

We conducted multivariable MR analysis to estimate the direct effects of education and general cognitive ability on our three smoking phenotypes. Multivariable MR estimates the effect of educational attainment on smoking conditional on general cognitive ability and the effect of general cognitive ability on smoking conditional on educational attainment. Multivariable MR includes both education and general cognitive ability as exposure variables in an MR regression with both of these exposures being predicted by a set of SNPs. This analysis controls for any correlation between education and general cognitive ability, and for any pleiotropic effect on smoking of the education SNPs through an effect of those SNPs on general cognitive ability and the general cognitive ability SNPs through an effect of those SNPs on education. Fig. 1 illustrates the model being estimated by our MR analyses, it is not necessary to specify for this analysis whether Fig. 1A or Fig. 1B represents the true model. The effects estimated are *β*_1_, the direct effect of education on smoking, and *β*_2_ the direct effect of general cognitive ability on smoking.

Multivariable MR is conducted by regressing each of the exposure variables, education and general cognitive ability, on all of the genetic instruments and all of the control variables to generate predicted values for each exposure that are not correlated with any potential confounders. The outcome of interest, smoking, is then regressed on these new predicted exposure variables and the control variables in a multivariable regression to obtain consistent estimates of the direct effect of each of education and general cognitive ability on smoking behaviour. As the instruments are not associated with any potential confounders this method controls for potential confounding of the relationship between the exposures, educational attainment and cognitive ability, and the outcome, smoking in the same way as in univariable MR. Multivariable MR is explained in more detail elsewhere.^9^ Multivariable MR uses values of educational attainment and general cognitive ability predicted from a set of SNPs which have been found to be associated with educational attainment or general cognitive ability in an independent sample. The inclusion of both exposures in the estimation does not introduce collider bias as the estimation is based on values of education and general cognitive ability predicted from SNPs that cannot depend on smoking behaviour, this is explained in more detail elsewhere. ^9^

Throughout the analysis we adjusted for age, sex, year of birth, and a year of birth × sex interaction term, in order to adjust for any changes in patterns of smoking behaviour over time. We also controlled for the top 10 genetic principal components to account for any residual population stratification.

To test the strength of the association of the instruments with the exposures we calculate the Sanderson-Windmeijer conditional F-statistic. This tests the ability of the SNPs to uniquely predict both education and cognitive ability. This is important because the results obtained from the multivariable MR analysis will be subject to weak instrument bias if either of education or cognitive ability are weakly predicted by the instruments. A value greater than 10 suggests that the instruments do predict the exposure well and the estimation result is not likely to be subject to weak instrument bias. ^9,39^

### Summary-level analysis

We additionally conducted univariable and multivariable two-sample summary data MR analysis of educational attainment and general cognitive ability on smoking initiation and cessation using summary data from genome-wide association studies (GWAS) of each of educational attainment, general cognitive ability and smoking behaviour.^9,40,41^ Ethical approval for each GWAS was obtained by the individual studies. This analysis only used publicly available summary data that does not include individual level data on any participants and so no further ethical approvals are necessary.

SNPs for educational attainment were taken from a large GWAS meta-analysis of 1.1 million individuals ^42^. We included all SNPs that were genome-wide significant (p<5×10-8) from this GWAS and pruned the results to exclude all SNPs with a pairwise R2 greater than 0.001. This gave 293 independent SNPs associated with educational attainment. For cognitive ability we used all SNPs which were genome wide significant in a GWAS of cognitive ability, estimated in a subset of the studies used in the educational attainment GWAS. These SNPs where also pruned to exclude SNPs with a pairwise R2 greater than 0.001 giving 100 independent SNPs. The effects of each SNP on smoking initiation and cessation was taken from a GWAS meta-analysis of 74,053 individuals smoking behaviour ^43^. There was limited overlap, of a maximum of 3,438 individuals from two studies, between the GWAS for educational attainment and smoking behaviour. This overlap represents 0.3% of the individuals in the educational attainment GWAS and 4.6% of the individuals in the smoking GWAS. There was no overlap between the GWAS used for cognitive ability and for smoking. For the univariable analysis we calculate the IVW estimate individually for each of educational attainment and general cognitive ability on smoking initiation and cessation.

For the multivariable MR analysis we included all SNPs which were genome wide significant in either the GWAS of education attainment or cognitive ability. This combined list was again pruned to exclude any SNPs with a pairwise R2 greater than 0.001 leaving 327 independent SNPs for the analysis. For summary data multivariable MR analysis the IVW framework can be extended by estimating the regression;

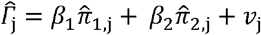

Using summary data estimates of the association between each SNP *j* (out of *L* where *L* is the total number of SNPs included in the analysis) and: smoking behaviour, 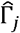; educational attainment, 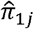; and cognitive ability, 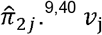.^9,40^ *ν_j_* is a random regression error term. All summary data MR analysis was conducted using MR base ^44^.

## Data Availability

Individual level analysis was conducted using UK Biobank (https://www.ukbiobank.ac.uk/). Access to the data is available to researchers through application to UK Biobank. Summary level analysis was conducted using publicly available data.^42,43^ The summary data for educational attainment and cognitive ability is available at https://www.thessgac.org/data, the summary data on smoking behaviour is available through MR base http://www.mrbase.org/.

## Code Availability

The code used to conduct all of the analysis reported is available on request from the corresponding author.

## Acknowledgements

This research has been conducted using the UK Biobank Resource as part of application 8786, and was supported by the UK Medical Research Council Integrative Epidemiology Unit at the University of Bristol (MM_UU_00011/1, MM_UU_00011/2, MM_UU_00011/7). MRM is a member of the UK Centre for Tobacco and Alcohol Studies, a UKCRC Public Health Research: Centre of Excellence. Funding from British Heart Foundation, Cancer Research UK, Economic and Social Research Council, Medical Research Council, and the National Institute for Health Research, under the auspices of the UK Clinical Research Collaboration, is gratefully acknowledged.

## Author Contributions

ES, GDS and MRM conceived the study. ES conducted the analyses. ES and MRM drafted the manuscript. All authors reviewed and approved the final version.

## References

1 Collaborators, G. B. D. T. Smoking prevalence and attributable disease burden in 195 countries and territories, 1990-2015: a systematic analysis from the Global Burden of Disease Study 2015. Lancet 389, 1885–1906, doi:10.1016/S0140-6736(17)30819-X (2017).

2 Davey Smith, G. & Ebrahim, S. ‘Mendelian randomization’: can genetic epidemiology contribute to understanding environmental determinants of disease? International journal of epidemiology 32, 1–22 (2003).

3 Tillmann, T. et al. Education and coronary heart disease: mendelian randomisation study. Bmj 358, j3542, doi:10.1136/bmj.j3542 (2017).

4 Gage, S. H., Davey Smith, G., Bowden, J. & Munafò, M. R. Investigating causality in associations between education and smoking: a two-sample Mendelian randomization study. International Journal of Epidemiology 47, 1131–1140, doi:10.1093/ije/dyy131 (2018).

5 Carter, A. R. et al. What explains the effect of education on cardiovascular disease? Applying Mendelian randomization to identify the consequences of education inequality. preprint avaliable at https://www.biorxiv.org/content/10.1101/488254v2, 488254, doi:10.1101/488254 (2018).

6 Deary, I. J., Strand, S., Smith, P. & Fernandes, C. Intelligence and educational achievement. Intelligence 35, 13–21, doi:https://doi.org/10.1016/j.intell.2006.02.001 (2007).

7 Deary, I. J. & Johnson, W. Intelligence and education: causal perceptions drive analytic processes and therefore conclusions. International Journal of Epidemiology 39, 1362–1369, doi:10.1093/ije/dyq072 (2010).

8 Strenze, T. Intelligence and socioeconomic success: A meta-analytic review of longitudinal research. Intelligence 35, 401–426, doi:https://doi.org/10.1016/j.intell.2006.09.004 (2007).

9 Sanderson, E., Davey Smith, G., Windmeijer, F. & Bowden, J. An examination of multivariable Mendelian randomization in the single sample and two-sample summary data settings. International Journal of Epidemiology, doi:10.1101/306209 (In press).

10 Fry, A. et al. Comparison of sociodemographic and health-related characteristics of UK Biobank participants with those of the general population. American journal of epidemiology 186, 1026–1034 (2017).

11 Hughes, R. A., Davies, N. M., Smith, G. D. & Tilling, K. Selection bias in instrumental variable analyses. Epidemiology, in press, 192237 (2018).

12 Munafò, M. & Smith, G. Biased estimates in mendelian randomization studies conducted in unrepresentative samples. JAMA Cardiology 3, 181–181, doi:10.1001/jamacardio.2017.4279 (2018).

13 Davey Smith, G. & Hemani, G. Mendelian randomization: genetic anchors for causal inference in epidemiological studies. Human Molecular Genetics 23, R89–R98, doi:10.1093/hmg/ddu328 (2014).

14 Bowden, J., Hemani, G. & Davey Smith, G. Detecting individual and global horizontal pleiotropy in Mendelian randomization: a job for the humble heterogeneity statistic? American Journal of Epidemiology, kwy185–kwy185, doi:10.1093/aje/kwy185 (2018).

15 Rees, J. M. B., Wood, A. M. & Burgess, S. Extending the MR-Egger method for multivariable Mendelian randomization to correct for both measured and unmeasured pleiotropy. Statistics in medicine 36, 4705–4718, doi:10.1002/sim.7492 (2017).

16 Davies, N. M., Dickson, M., Davey Smith, G., van den Berg, G. J. & Windmeijer, F. The causal effects of education on health outcomes in the UK Biobank. Nature Human Behaviour 2, 117–125, doi:10.1038/s41562-017-0279-y (2018).

17 Daly, M. & Egan, M. Childhood cognitive ability and smoking initiation, relapse and cessation throughout adulthood: evidence from two British cohort studies. Addiction (Abingdon, England) 112, 651–659, doi:10.1111/add.13554 (2017).

18 Davies, L. E. M., Kuipers, M. A. G., Junger, M. & Kunst, A. E. The role of self-control and cognitive functioning in educational inequalities in adolescent smoking and binge drinking. BMC Public Health 17, 714, doi:10.1186/s12889-017-4753-2 (2017).

19 Hemani, G., Bowden, J. & Davey Smith, G. Evaluating the potential role of pleiotropy in Mendelian randomization studies. Human Molecular Genetics 27, R195–R208, doi:10.1093/hmg/ddy163 (2018).

20 Kong, A. et al. The nature of nurture: Effects of parental genotypes. Science 359, 424–428 (2018).

21 Conti, G., Heckman, J. & Urzua, S. The Education-Health Gradient. American Economic Review 100, 234–238, doi:10.1257/aer.100.2.234 (2010).

22 Næss, Ø., Hoff, D. A., Lawlor, D. & Mortensen, L. H. Education and adult cause-specific mortality—examining the impact of family factors shared by 871 367 Norwegian siblings. International Journal of Epidemiology 41, 1683–1691, doi:10.1093/ije/dys143 (2012).

23 Lawlor, D. A., Tilling, K. & Davey Smith, G. Triangulation in aetiological epidemiology. International Journal of Epidemiology 45, 1866–1886 (2016).

24 Munafo, M. R., Tilling, K., Taylor, A. E., Evans, D. M. & Davey Smith, G. Collider scope: when selection bias can substantially influence observed associations. International journal of epidemiology, doi:10.1093/ije/dyx206 (2017).

25 Allen, N. E., Sudlow, C., Peakman, T. & Collins, R. UK biobank data: come and get it. Science Translational Medicine (2014).

26 Collins, R. What makes UK Biobank special? The Lancet 379, 1173–1174 (2012).

27 Bycroft, C. et al. Genome-wide genetic data on~ 500,000 UK Biobank participants. preprint avaliable at https://www.biorxiv.org/content/10.1101/166298v1, 166298 (2017).

28 Manichaikul, A. et al. Robust relationship inference in genome-wide association studies. Bioinformatics 26, 2867–2873 (2010).

29 Mitchell, R., Hemani, G., Dudding, T. & Paternoster, L. UK Biobank Genetic Data: MRC-IEU Quality Control, Version 1. University of Bristol (2017).

30 Hagenaars, S. P. et al. Shared genetic aetiology between cognitive functions and physical and mental health in UK Biobank (N=112□151) and 24 GWAS consortia. Molecular Psychiatry 21, 1624 (2016).

31 Sniekers, S. et al. Genome-wide association meta-analysis of 78,308 individuals identifies new loci and genes influencing human intelligence. Nature Genetics (2017).

32 Deary, I. J., Penke, L. & Johnson, W. The neuroscience of human intelligence differences. Nature Reviews Neuroscience 11, 201, doi:10.1038/nrn2793 (2010).

33 Okbay, A. et al. Genome-wide association study identifies 74 loci associated with educational attainment. Nature 533, 539–542, doi:10.1038/nature17671 (2016).

34 Sniekers, S. et al. Genome-wide association meta-analysis of 78,308 individuals identifies new loci and genes influencing human intelligence. Nature genetics 49, 1107–1112, doi:10.1038/ng.3869 (2017).

35 Taylor, A. E. et al. Mendelian randomization in health research: using appropriate genetic variants and avoiding biased estimates. Economics & Human Biology 13, 99–106 (2014).

36 Burgess, S. & Thompson, S. G. Use of allele scores as instrumental variables for Mendelian randomization. Int. J. Epidemiol. 42, 1134–1144, doi:10.1093/ije/dyt093 (2013).

37 Davey Smith, G. & Ebrahim, S. ‘Mendelian randomization’: can genetic epidemiology contribute to understanding environmental determinants of disease? International journal of epidemiology 32, 1–22 (2003).

38 Lawlor, D. A., Harbord, R. M., Sterne, J. A., Timpson, N. & Davey Smith, G. Mendelian randomization: using genes as instruments for making causal inferences in epidemiology. Statistics in medicine 27, 1133–1163 (2008).

39 Sanderson, E. & Windmeijer, F. A weak instrument F-test in linear IV models with multiple endogenous variables. Journal of Econometrics 190, 212–221 (2016).

40 Burgess, S. & Thompson, S. G. Multivariable Mendelian Randomization: The Use of Pleiotropic Genetic Variants to Estimate Causal Effects. American Journal of Epidemiology 181, 251–260, doi:10.1093/aje/kwu283 (2015).

41 Burgess, S., Dudbridge, F. & Thompson, S. G. Re:“Multivariable Mendelian randomization: the use of pleiotropic genetic variants to estimate causal effects”. American journal of epidemiology 181, 290–291 (2015).

42 Lee, J. J. et al. Gene discovery and polygenic prediction from a genome-wide association study of educational attainment in 1.1 million individuals. Nature genetics 50, 1112 (2018).

43 Furberg, H. et al. Genome-wide meta-analyses identify multiple loci associated with smoking behavior. Nature genetics 42, 441 (2010).

44 Hemani, G. et al. The MR-Base platform supports systematic causal inference across the human phenome. eLife 7, e34408 (2018).

