## Supplementary Material for "Mendelian randomisation analysis of the effect of educational attainment and cognitive ability on smoking behaviour"

**Supplementary Figure 1**

a.
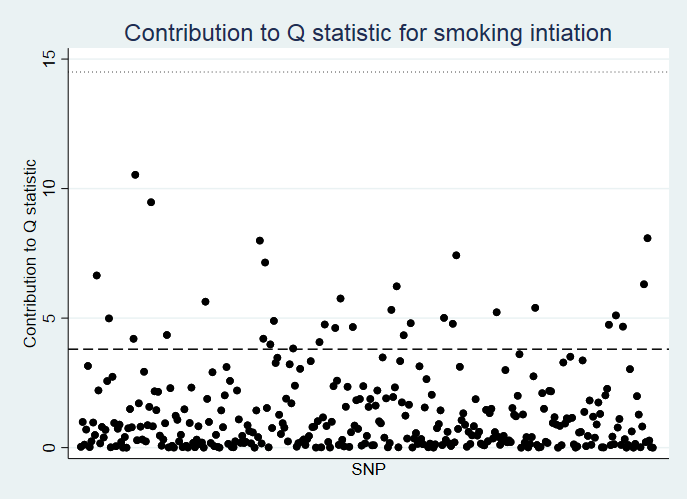

b.
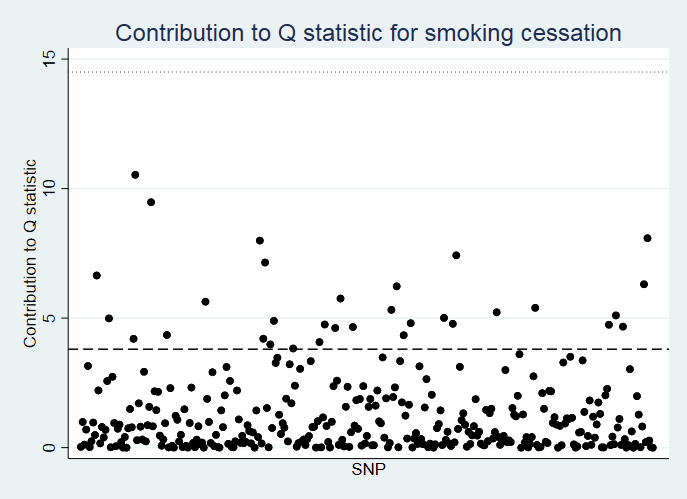

Contribution to the overall Q statistic for heterogeneity in the summary-data pleiotropy test of each individual SNP. Figure S.1a. gives the contribution of each SNP to the Q-statistic for smoking initiation. Figure S.1b. gives the contribution of each SNP to the Q-statistic for smoking cessation.

**Supplementary Table 1 – Distribution of self-reported age left school for highest reported educational qualification.**

| Self-reported age | None | CSE / O level / GCSE | NVQ / HND / HNC | A level | Other professional qualification * |
| --- | --- | --- | --- | --- | --- |
| *N* | ***14,348*** | ***32,056*** | ***7,055*** | ***14,839*** | ***5,973*** |
| <13 | 0.2 | 0.0 | 0.1 | 0.0 | 0.1 |
| 13 | 0.1 | 0.0 | 0.0 | 0.0 | 0.0 |
| 14 | 2.5 | 0.2 | 1.3 | 0.2 | 0.8 |
| 15 | 73.5 | 10.8 | 39.1 | 4.1 | 22.6 |
| 16 | 20.6 | 51.0 | 27.9 | 13.4 | 20.8 |
| 17 | 1.9 | 21.1 | 6.7 | 11.1 | 10.1 |
| 18 | 0.7 | 10.7 | 5.6 | 37.3 | 12.8 |
| 19 | 0.1 | 1.7 | 1.7 | 10.1 | 2.4 |
| 20 | 0.1 | 1.1 | 2.4 | 5.9 | 3.2 |
| 21 | 0.2 | 1.8 | 5.9 | 12.0 | 17.7 |
| 22 | 0.1 | 0.4 | 2.9 | 3.1 | 4.6 |
| 23 | 0.0 | 0.2 | 1.9 | 1.2 | 1.6 |
| 24 | 0.0 | 0.1 | 1.0 | 0.5 | 0.8 |
| 25 | 0.0 | 0.2 | 1.1 | 0.4 | 0.7 |
| >25 | 0.0 | 0.4 | 2.4 | 0.8 | 1.7 |

The proportion of individuals who report each age for leaving school for each level of qualification. Individuals who reported having a degree were not asked their age at leaving full time education and therefore are excluded from this table.

**Supplementary Table 2. Multivariable MR estimates of the effect of education and general cognitive ability on smoking behaviour using self-reported age at leaving education. Risk difference estimates from individual level data.**

|  | *Current Smoking* | | *Smoking Initiation* | | *Smoking cessation* | |
| --- | --- | --- | --- | --- | --- | --- |
|  | **Single variable MR** | **Multivariable MR** | **Single variable MR** | **Multivariable MR** | **Single variable MR** | **Multivariable MR** |
| *Age completed Education* | | |  |  |  |  |
| Effect | -0.025 | -0.038 | -0.052 | -0.085 | 0.035 | 0.049 |
| SE | 0.005 | 0.011 | 0.008 | 0.021 | 0.009 | 0.020 |
| 95% CI | -0.034 to -0.016 | -0.060, -0.016 | -0.068 to -0.035 | -0.126, -0.044 | 0.017 to 0.053 | 0.009, 0.088 |
| P-value | <0.001 | 0.001 | <0.001 | <0.001 | <0.001 | 0.017 |
| F | 513.46 | 453.84 | 513.46 | 453.84 | 237.68 | 196.63 |
| Conditional F | - | 78.60 | - | 78.60 | - | 46.20 |
| *Cognitive ability score* | | |  |  |  |  |
| Effect | -0.026 | 0.047 | -0.043 | 0.122 | 0.036 | -0.052 |
| SE | 0.009 | 0.027 | 0.017 | 0.051 | 0.019 | 0.048 |
| 95% CI | -0.044 to -0.008 | -0.006, 0.100 | -0.076 to -0.010 | 0.023, 0.221 | -0.0004 to 0.073 | -0.147 to 0.043 |
| P-value | 0.004 | 0.082 | 0.011 | 0.016 | 0.053 | 0.285 |
| F | 920.27 | 556.69 | 920.27 | 556.69 | 429.70 | 258.26 |
| Conditional F | - | 79.98 | - | 79.98 | - | 47.08 |
| Sample Size | 119,260 | | 119,260 | | 52,228 | |

Estimates based on self-reported age at leaving education. Estimates of the effect of education and general cognitive ability on smoking behaviour from single and multivariable mendelian randomisation using individual-level data. All regressions also include a full set of adjustments: age, sex, year of birth and gender interacted with year of birth. Mendelian randomisation regressions also control for 10 genetic principal components. Non-European and related individuals have been excluded from the analysis.

**Supplementary Table 3. Multivariable MR estimates of the effect of education and general cognitive ability on smoking behaviour using measure of cognitive ability taken from the clinic only. Risk difference estimates from individual level data.**

|  | *Current Smoking* | | *Smoking Initiation* | | *Smoking cessation* | |
| --- | --- | --- | --- | --- | --- | --- |
|  | **Single variable MR** | **Multivariable MR** | **Single variable MR** | **Multivariable MR** | **Single variable MR** | **Multivariable MR** |
| *Age completed Education* | | |  |  |  |  |
| Effect | -0.029 | -0.042 | -0.045 | -0.084 | 0.044 | 0.057 |
| SE | 0.006 | 0.017 | 0.011 | 0.031 | 0.013 | 0.031 |
| 95% CI | -0.041 to -0.017 | -0.075 to -0.008 | -0.067 to -0.023 | -0.144 to -0.024 | 0.020 to 0.069 | -0.003 to 0.118 |
| P-value | <0.001 | 0.014 | <0.001 | 0.006 | <0.001 | 0.064 |
| F | 368.38 | 360.46 | 368.38 | 360.46 | 175.59 | 159.94 |
| Conditional F | - | 46.92 | - | 46.92 | - | 26.95 |
| *Cognitive ability score* | | |  |  |  |  |
| Effect | -0.034 | 0.039 | -0.031 | 0.117 | 0.057 | -0.040 |
| SE | 0.010 | 0.036 | 0.018 | 0.064 | 0.021 | 0.064 |
| 95% CI | -0.054 to -0.015 | -0.031 to 0.108 | -0.066 to 0.005 | -0.009 to 0.243 | 0.015 to 0.098 | -0.166 to 0.086 |
| P-value | 0.001 | 0.272 | 0.093 | 0.069 | 0.007 | 0.535 |
| F | 797.60 | 488.80 | 797.60 | 488.80 | 355.42 | 216.48 |
| Conditional F | - | 47.80 | - | 47.80 | - | 27.61 |
| Sample Size | 85,206 | | 85,206 | | 37,935 | |

Estimated effects using cognitive ability scores from the clinic only. Estimates of the effect of education and general cognitive ability on smoking behaviour from single and multivariable mendelian randomisation using individual-level data. All regressions also include a full set of adjustments: age, sex, year of birth and gender interacted with year of birth. Mendelian randomisation regressions also control for 10 genetic principal components. Non-European and related individuals have been excluded from the analysis.

**Supplementary Table 4. Multivariable MR estimates of the effect of education and general cognitive ability on smoking behaviour estimated using weighting. Risk difference estimates from individual level data.**

|  | *Current Smoking* | | | *Smoking Initiation* | | *Smoking cessation* | |
| --- | --- | --- | --- | --- | --- | --- | --- |
|  | **Single variable MR** | **Multivariable MR** | **Single variable MR** | | **Multivariable MR** | **Single variable MR** | **Multivariable MR** |
| *Age completed Education* | | | |  |  |  |  |
| Effect | -0.030 | -0.047 | | -0.057 | -0.101 | 0.043 | 0.060 |
| SE | 0.005 | 0.013 | | 0.009 | 0.024 | 0.011 | 0.025 |
| 95% CI | -0.041 to -0.020 | -0.074 to -0.021 | | -0.074 to -0.039 | -0.147, -0.055 | 0.022 to 0.064 | 0.012, 0.109 |
| P-value | <0.001 | <0.001 | | <0.001 | <0.001 | <0.001 | 0.015 |
| F | 589.96 | 518.34 | | 589.96 | 518.34 | 269.14 | 222.36 |
| Conditional F | - | 82.61 | | - | 82.61 | - | 45.80 |
| *Cognitive ability score* | | | |  |  |  |  |
| Effect | -0.032 | 0.058 | | -0.042 | 0.150 | 0.047 | -0.058 |
| SE | 0.010 | 0.031 | | 0.018 | 0.056 | 0.020 | 0.055 |
| 95% CI | -0.052 to -0.012 | -0.004 to 0.120 | | -0.077 to -0.007 | 0.041, 0.260 | -0.007 to 0.087 | -0.167, 0.051 |
| P-value | 0.002 | 0.065 | | 0.018 | 0.007 | 0.022 | 0.299 |
| F | 836.03 | 522.38 | |  | 522.38 | 374.07 | 232.75 |
| Conditional F | - | 82.54 | | - | 82.54 | - | 45.85 |
| Sample Size | 120,050 | | | 120,050 | | 52,605 | |

Estimated effects using a weighting calculated to adjust the sample based on the estimated proportion of individuals of the same age range in the UK population who have no qualifications or degree level qualifications, based on 2011 UK census figures. Estimates of the effect of education and general cognitive ability on smoking behaviour from single and multivariable mendelian randomisation using individual-level data. All regressions also include a full set of adjustments: age, sex, year of birth and gender interacted with year of birth. Mendelian randomisation regressions also control for 10 genetic principal components. Non-European and related individuals have been excluded from the analysis.

**Supplementary Table 5. Estimated Q statistics for two-sample MVMR analysis, with and without potentially pleiotropic outlying SNPs.**

|  |  | Smoking Initiation | Smoking Cessation |
| --- | --- | --- | --- |
| Including All SNPs | Q-statistic | 445 | 356 |
|  | P-value | <0.001 | 0.122 |
|  | No. SNPs | 327 | 327 |
| Potentially Pleiotropic SNPs removed | Q-statistic | 266 | 251 |
|  | P-value | 0.878 | 0.993 |
|  | No. SNPs | 295 | 310 |

**Supplementary Table 6. Estimates of the effect of educational attainment and general cognitive ability with potentially pleiotropic SNPs excluded. Odds ratio estimates from a two-sample MR analysis**

|  | Smoking Initiation | | Smoking cessation | |
| --- | --- | --- | --- | --- |
|  | **Single variable MR** | **Multivariable MR** | **Single variable MR** | **Multivariable MR** |
| *Age completed Education* | | |  |  |
| Effect | 0.610 | 0.620 | 1.848 | 2.457 |
| 95% CI | 0.559 to 0.715 | 0.449 to 0.770 | 1.582 to 2.158 | 1.866 to 3.238 |
| P-value | <0.001 | <0.001 | <0.001 | <0.001 |
| *Cognitive ability score* | | |  |  |
| Effect | 0.687 | 1.023 | 1.490 | 0.709 |
| 95% CI | 0.604 to 0.780 | 0.822 to 1.275 | 1.266 to 1.704 | 0.539 to 0.933 |
| P-value | <0.001 | 0.834 | <0.001 | 0.014 |

Estimates of the effect of education and general cognitive ability on smoking behaviour from single and multivariable mendelian randomisation using summary-level data. SNPs with a statistically significant contribution to the Q-statistic to detect potential pleiotropy (p<0.05) have been excluded from the analysis. Effects given are odds ratios.

Total number of SNPs included; smoking initiation – 295, smoking cessation – 310.

Summary data statistics obtained from publicly available results from GWAS studies on educational attainment, cognitive ability (1) and smoking behaviour (2).

**Supplementary Table 7. Results from multivariable MR Egger estimates of the effect of education and general cognitive ability on smoking behaviour.**

|  | *Smoking Initiation* | | | *Smoking Cessation* | | |
| --- | --- | --- | --- | --- | --- | --- |
|  | **Single variable MR** | | **Multivariable MR** | **Single variable MR** | | **Multivariable MR** |
| *Age completed education* | | | |  |  |  |
| Effect | 0.847 |  | 0.605 | 1.887 |  | 2.617 |
| 95% CI | 0.520 to 1.387 |  | 0.468, 0.783 | 1.029 to 3.459 |  | 1.907, 3.586 |
| P-value | 0.513 |  | <0.001 | 0.040 |  | <0.001 |
| *Cognitive ability score* | | | |  |  |  |
| Effect |  | 1.046 | 1.110 |  | 0.872 | 0.686 |
| 95% CI |  | 0.797 to 1.373 | 0.856, 1.436 |  | 0.619 to 1.226 | 0.501, 0.942 |
| P-value |  | 0.744 | 0.431 |  | 0.429 | 0.020 |
| *Constant (Estimated pleiotropic effect)* | | | |  |  |  |
| Effect | 0.997 | 0.995 | 1.001 | 1.000 | 1.007 | 1.000 |
| 95% CI | 0.991 to 1.003 | 0.992 to 0.998 | 0.999, 1.003 | 0.993 to 1.007 | 1.003 to 1.012 | 0.998, 1.002 |
| P-value | 0.285 | 0.002 | 0.306 | 0.957 | <0.001 | 0.959 |

Multivariable MR Egger (3) estimates of the effect of educational attainment and cognitive ability on smoking behaviour. Effects of SNPs on each of the exposures and the outcome are taken from GWAS results (1, 2) Effects given are odds ratios.

For MR Egger analysis it is important to align all of the SNPs so they have a positive effect on the exposure; it is not clear how to do this in multivariable MR Egger and therefore we repeated the analysis with all of the effects aligned in the positive direction for firstly the effect on education and then their effect on cognitive ability. This adjustment made no difference to the results obtained.

**Supplementary Table 8 – Power calculations for individual level and between sibling analysis.**

|  | *Effect Size* | | | | | | | |
| --- | --- | --- | --- | --- | --- | --- | --- | --- |
|  | **0.01** | **0.025** | **0.05** | **0.075** | **0.1** | **0.15** | **0.2** | **0.25** |
| Individual analysis | 1.4% | 4.5% | 19.6% | 36.1% | 54.6% | 76.6% | 85.7% | 91.0% |
| Between sibling analysis | 0.6% | 2.4% | 9.1% | 25.5% | 42.1% | 62.2% | 76.9% | 85.7% |

Power to reject the null hypothesis of the effect of educational attainment on smoking in a multivariable MR study for varying effect sizes. Simulations are set up with N = 11,448, grouped into 5,724 sibling pairs with the correlations in the simulated data set to reflect those observed in the UK Biobank data. 1,000 repetitions per simulation

**References**

1.	Lee JJ, Wedow R, Okbay A, Kong E, Maghzian O, Zacher M, et al. Gene discovery and polygenic prediction from a genome-wide association study of educational attainment in 1.1 million individuals. Nature genetics. 2018;50(8):1112.

2.	Furberg H, Kim Y, Dackor J, Boerwinkle E, Franceschini N, Ardissino D, et al. Genome-wide meta-analyses identify multiple loci associated with smoking behavior. Nature genetics. 2010;42(5):441.

3.	Rees JMB, Wood AM, Burgess S. Extending the MR-Egger method for multivariable Mendelian randomization to correct for both measured and unmeasured pleiotropy. Statistics in medicine. 2017;36(29):4705-18.
